# Identification of a crucial *INO2* allele for enhancing ethanol resistance in an industrial fermentation strain of *Saccharomyces cerevisiae*

**DOI:** 10.1101/2024.09.23.614527

**Authors:** Sonia Albillos-Arenal, Javier Alonso del Real, María Lairón-Peris, Eladio Barrio, Amparo Querol

## Abstract

Ethanol toxicity is a major challenge for *S. cerevisiae* during fermentation, affecting its growth and influencing the process. This study investigated the molecular mechanisms of ethanol tolerance using transcriptomic analysis of three *S. cerevisiae* strains with varying ethanol resistance. We identified distinct responses in membrane lipid synthesis genes, particularly in ergosterol biosynthesis, regulated by the Ino2p transcription factor. A variant of Ino2p with V263I and H86R amino acid replacements was exclusive to ethanol-tolerant strains. CRISPR-Cas9-mediated point mutations in the *INO2* gene of the highly tolerant strain AJ4 resulted in decreased ethanol tolerance. Our findings demonstrate the crucial role of Ino2p in ethanol tolerance through regulation of lipid synthesis and membrane composition, highlighting the complex interplay of trans elements in strain-specific ethanol resistance

**IMPORTANCE:** This study provides critical insights into the molecular basis of ethanol tolerance in *S. cerevisiae*, a key trait for improving industrial fermentation processes. By identifying specific genetic variants in the Ino2p transcription factor and their impact on ethanol resistance, we reveal potential targets for enhancing yeast strain performance in high-ethanol environments. Our findings not only contribute to the fundamental understanding of stress response mechanisms in yeast but also offer practical implications for strain engineering in the biotechnology and beverage industries. The unexpected magnitude of the Ino2p variants’ effect on ethanol tolerance underscores the importance of considering strain-specific genetic backgrounds in metabolic engineering strategies

## INTRODUCTION

During fermentation, yeast cells are dynamically exposed various interrelated stresses, including osmotic (1), oxidative (2), thermal (3), ethanol (4), and starvation (5). These stress conditions can significantly affect the yeast population and fermentation efficiency (6). Yeast cells have evolved to be exceptionally capable of surviving these sudden and harsh changes. Among all the stresses, ethanol toxicity is a major hindrance to fermentation, impacting cell growth, cell cycle regulation, and various metabolic functions in microorganisms. Studies have shown that ethanol significantly influences the fermentation capabilities of industrial *Saccharomyces* yeast strains (7). This is particularly relevant for biotechnological applications, such as wine fermentation or bioethanol production. Consequently, stress tolerance mechanisms are crucial for the efficiency of yeast cell growth and metabolism (8). The challenges posed by these stresses have led to the development of robust strain platforms aimed at improving fermentation kinetics, product yield, and cellular robustness under process conditions (9). Despite these advancements, achieving optimal ethanol yields during fermentation remains a significant technical challenge. This underscores the critical need for continued research to further improve strain performance and process efficiency in ethanol production.

Abundant literature has shown how yeasts respond to ethanol stress through general stress response mechanisms (10–12). Yeast cells employ a multifaceted approach to respond to ethanol stress, including reprogramming cellular activities, protecting essential components, and promoting cell survival during stress conditions. They activate various protective mechanisms, such as accumulating heat shock proteins, adjusting membrane sterols, and increasing trehalose levels to stabilize proteins and membranes (12, 13). Additionally, yeasts adapt their intracellular glucose metabolism pathways, regulate fatty acid and amino acid metabolisms, and reconstruct cell membrane structures to mitigate the damage caused by ethanol (8). Arginine has been found to play a protective role in yeast cells against ethanol stress by maintaining the integrity of the cell wall and cytoplasmic membrane (14). The upregulation of heat shock protein and trehalose synthesis genes is a common response to various fermentation-associated stresses, including high osmolarity, increased ethanol concentration, the production of reactive oxygen species (ROS), and elevated temperature (15). While common stress response pathways, such as energy metabolism, amino acid metabolism, and fatty acid metabolism, are shared among yeast species, the specific genes and metabolic pathways involved in ethanol tolerance can vary between strains. A study using large-scale data integration revealed that long non-coding RNAs (lncRNAs) play a strain-specific role in the ethanol stress response (16). Additionally, these lncRNAs can modulate gene expression through cis-regulation of adjacent genes or trans-regulation of distant genes (17). Regulatory network studies and transcriptomic analyses reveal that cells prepare for stress relief by prioritizing the activation of life-essential systems, such as longevity pathways, peroxisomal, energy, lipid, and RNA/protein metabolisms. The differences in response to ethanol stress between high and low ethanol-tolerant phenotypes occur after cell signaling impacts the longevity and peroxisomal pathways, with catalase (*CTA1*) and reactive oxygen species playing critical roles. The study also emphasizes the significance of specific lipid metabolism pathways, as well as the roles of degradation processes and membrane structures in enhancing ethanol tolerance.

The fatty acid composition of lipid membranes has been linked to ethanol tolerance in various *Saccharomyces* strains (18–22). However, the precise mechanisms underlying these associations require further elucidation, particularly concerning gene expression changes in lipid biosynthesis.

Transcriptomics analyses have become popular tools for studying gene expression levels in specific biological contexts. Next-generation sequencing technologies, such as RNA sequencing (RNAseq), have revolutionized transcriptome-wide analysis, enabling the comparison of gene expression between different conditions and strains (16)

This study aimed to delve deeper into the molecular mechanisms and pathways responsible for varying ethanol tolerances in three *S. cerevisiae* strains. We hypothesize that these tolerance mechanism are primarily related to membrane lipid composition and the regulation of genes involved in lipid synthesis. What sets this study apart is the use of previously characterized *S. cerevisiae* strains (MY3, MY26, and AJ4) with known ethanol tolerance and membrane composition (21).

## RESULTS

### Transcriptomic analysis of *S. cerevisiae* strains with varying ethanol tolerance during their growth in ethanol media

To determine transcriptomic differences in response to ethanol, we selected three *S. cerevisiae* strains isolated which are highly, moderately, and slightly tolerant to ethanol, respectively: AJ4, MY3, and MY26 (21). The transcriptomes of the selected strains were analyzed at three time points, using the beginning of the fermentation, before ethanol addition, as a reference (*t0*). The first samples were taken at the early exponential phase (*t1*), the second at the late exponential phase (*t2*), and the last at the stationary phase (*t3*). The specific times at which each strain reached these growth stages under different ethanol conditions were determined based on their growth dynamics (Figure S1). The three strains showed similar growth when no ethanol was added. With 6% ethanol, AJ4 and MY26 grew similarly, but MY3 was noticeably affected. At 10% ethanol, all strains were severely affected, but AJ4 was the most tolerant, followed by MY3.

Transcriptomic analysis was performed taking *t0* as a control to identify differentially expressed (DE) genes among strains at each fermentation stage and ethanol concentration. Principal component analysis (PCA) of all samples showed that they primarily clustered by strains (**Figure 1**). Samples at t3 were more distinct, especially without ethanol. For t3 samples with 6% or 10% ethanol, the separation depended on the strain. Despite the high stress of 10% ethanol, these samples did not cluster separately, suggesting that strain and growth phase had a stronger impact on gene expression than ethanol concentration. As PCA analysis was based on the top 500 genes presenting more expression variability among all samples, it seems that the changes provoked by the variable “ethanol concentration” are masked by the ones by the variable “strain”, or even the variable “growth phase” (23).

**Figure 1.**
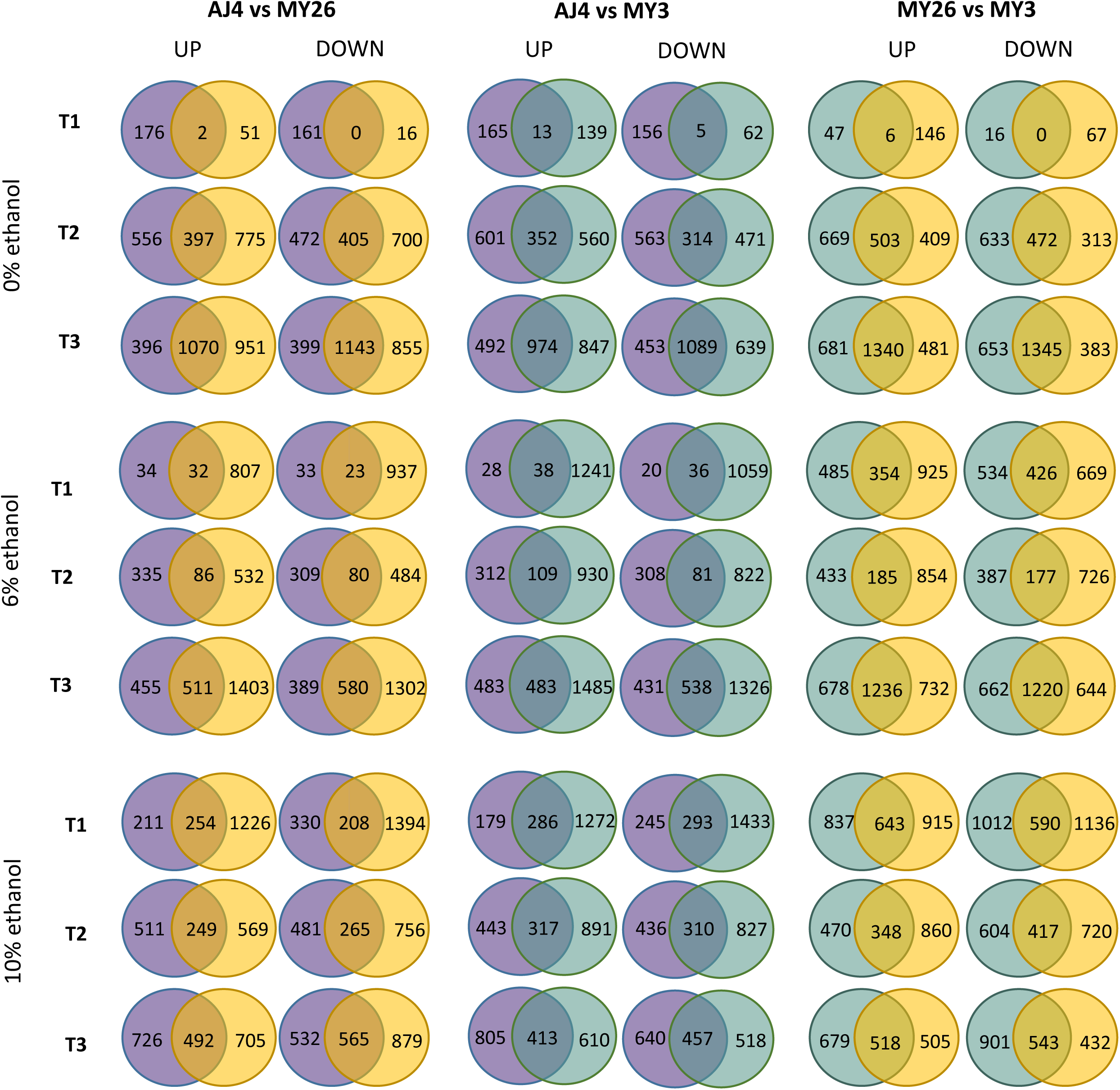
PCA analysis based on the top 500 genes with the highest expression variability across all samples. *S. cerevisiae* strains AJ4, MY3 and MY26 are represented by circles with purple, blue and yellow outlines, respectively. Ethanol (EtOH) concentrations of 0%, 6%, and 10% are indicated by blue, yellow, and red circle filling, respectively.

To focus on the ethanol effect in each strain, differential expression at every time point with 6% and 10% ethanol compared to 0% ethanol was assessed (Table S1). The number of DE genes increased over time, likely due to a decrease in the synchronization of the physiologic state of the cells. This was partly a consequence of the transcriptomic response to ethanol, which began when different ethanol concentrations were added, leading to a complex gene regulation cascade.

Starting with the analysis at *t1*, it is noteworthy that the stress-responsive genes *HSP26*, *HSP32,* and *SSA3* were strongly overexpressed (log_2_fold > 2.5) in all three strains under 10% ethanol stress. However under 10% ethanol at *t1*, strains MY3 and MY26 shared another set of highly overexpressed genes related to sporulation and mating: *ADY2, GAS4, HO, MAM1, MND1, PRM1, SGA1, SHC1, SPO13, SPS1,* and *SPS100.* This suggests that less tolerant strains prepare to produce spores to endure highly stressful conditions for long periods, while AJ4 can cope without such preparation. In contrast, under 6% ethanol at *t1*, the only gene commonly overexpressed among the three strains was *FMP45.* In this condition, the stress-responsive genes *DDR2* and *HSP12* were overexpressed in the two most sensitive strains MY3 and MY26. For AJ4, the overexpressed genes included *CTR1, CTR3, FET3, FIT2*, *FRE1, HSP26, HSP30,* and *SSA4.* At 10% ethanol *t1*, among the strongly repressed genes (log_2_fold < -2.5), the sensitive strains MY3 and MY26 shared more repressed genes with each other than with AJ4. These genes included *ADE1, ADE2, ADE13, ADE17, COB, COX1, COX2,* and *COX3*.

At *t2*, under 10% ethanol, few genes were strongly overexpressed in all three strains, but the stress response genes *HSP12* and *DDR2* were consistently found. Among the repressed genes at *t2* under 10% ethanol, *AGP1, DUR1,2, GAP1,* and *MEP2* were repressed in all strains. Additionally, strains MY3 and MY26 had *GAL2* and *HXT2* repressed.

At *t3*, the unspecific hexose transporters HXT3 and HXT4 were overexpressed in all strains. *ELO3* was also overexpressed in the three strains, whilst *ELO2* was exclusively overexpressed in the sensitive strains MY3 and MY26. A notable response at *t3* under 10% ethanol was the downregulation of aerobic respiration and fatty acid beta-oxidation in all strains, especially in MY3 and MY26.

A GO-term enrichment analysis (Table S2) showed some common aspects among the three strains, including significant rewiring of nitrogen and carbon metabolisms, and repression of genes involved in lipid synthesis and membrane regulation pathways, such as the tricarboxylic acid cycle and ergosterol synthesis. Stress response and protein refolding were also prominent.

Thus, we observed that the response to ethanol involves elements of the general stress response, which have varying consequences depending on the strain. Plasma membrane regulation could play an important role in this response.

### Importance of plasma membrane and lipid metabolism in strain-specific response to ethanol

To examine how gene expression varies at different stages of growth stages in comparison to a control situation for each strain, we established *t0* as the reference point in our experimental design. Specifically, we analyzed differentially expressed (DE) genes at each time point and ethanol condition by comparing each strain to the other two. This analysis involved identifying the intersection of DE gene sets. The large number of genes exclusive to each strain indicates highly variable transcriptome profiles among the studied strains (**Figure 2)**. Detailed lists of DE genes for strain-specific genes and the functional GO-term enrichment analyses derived from them can be found in the supplementary material (Tables S3-S5).

**Figure 2.**
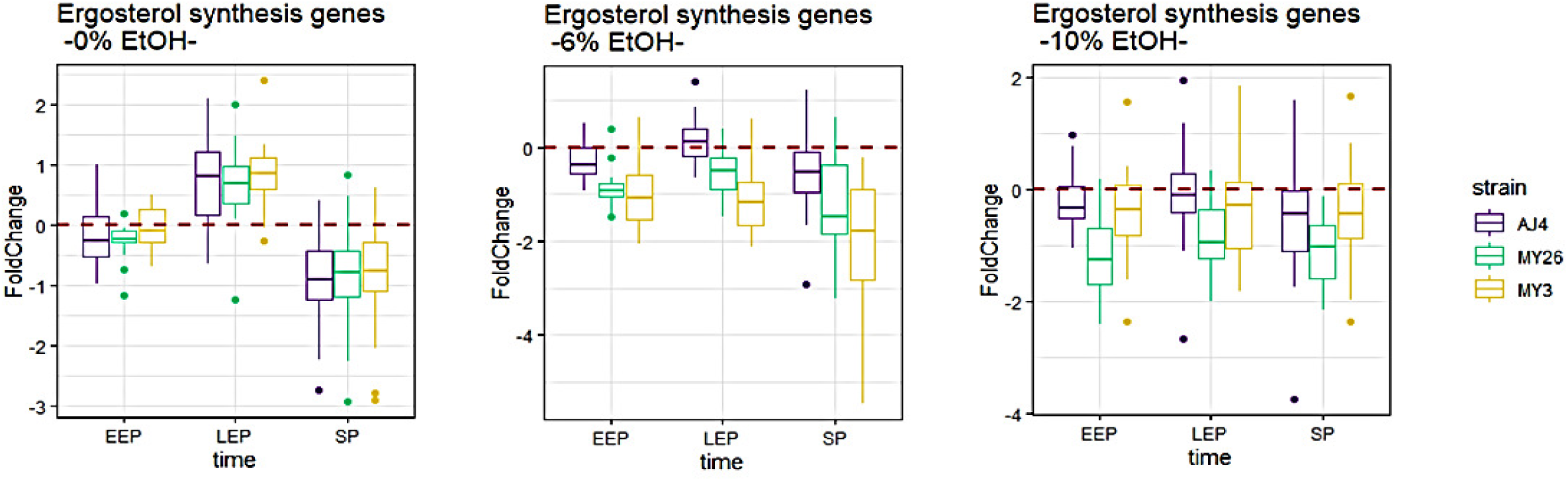
A Venn diagram illustrates the genes uniquely upregulated and downregulated in *S. cerevisiae* strains AJ4, MY3, and MY26 under three ethanol concentrations, at three time points. The strains were grown in GPY media containing 0%, 6%, and 10% ethanol, with samples collected at times *t1*, *t2*, and *t3*, and compared to *t0*. Differentially expressed genes were identified through DE analysis, applying a threshold for an adjusted p-value < 0.05 (Benjamini-Hochberg correction).

Due to the large input sample sizes for enrichment analyses and the application of Bonferroni corrections, some cellular functions with DE genes may be masked in our results. Many of the identified categories are directly related to general stress response, cell division, ribosomal translation and central carbon metabolism. These functions are integral to stress response and growth impairment, which were anticipated under the experimental conditions. The more sensitive strains, MY3 and MY26, exhibited many of these functional categories from their exclusively DE genes in comparison to AJ4.

Interestingly, numerous other groups of GO terms were related to both plasma membrane elements and lipid biosynthesis. The “Plasma Membrane Enriched Fraction” term was obtained from AJ4 repressed genes that were not present in MY3 or MY26. However, terms like “membrane”, “integral to membrane”, and “plasma membrane” were also found for MY26 and MY3. Additionally, categories “cellular lipid metabolic process” or “lipid biosynthetic process” were present in all three strains. Delving into more specific metabolic pathways, the “phospholipid biosynthetic process” term was identified in MY26 upregulated genes, distinct from MY3 at *t1* with 6% ethanol. This may indicate plasma membrane adjustments. However, ergosterol biosynthesis was the most recurrent family of GO terms for MY3 and MY26 repressed genes compared to AJ4, appearing at both ethanol concentrations at different time points. While translation or cell cycle arrest can result from ethanol-induced stress, the plasma membrane acts as the first barrier against ethanol, suggesting that yeasts may adjust to ethanol toxicity through plasma membrane remodeling.

Consequently, genes related to lipid metabolism and membrane homeostasis were investigated in detail by filtering out all the genes not included in the functional categories “plasma membrane” (GO:0005886) or “lipid metabolism” (GO:0044255) from the Gene Ontology. Interestingly, under 6% and 10% ethanol conditions, many genes coding for enzymes involved in ergosterol biosynthesis were repressed at every time point in the sensitive strains MY26 and MY3, but not in AJ4. In contrast, the expression of these genes was similar for all strains in the absence of ethanol (**Figure 3**). Examining the expression of each gene in the pathway revealed that most of them were repressed in MY26 and MY3 under ethanol conditions, rather than just one or a few genes standing out.

**Figure 3.**
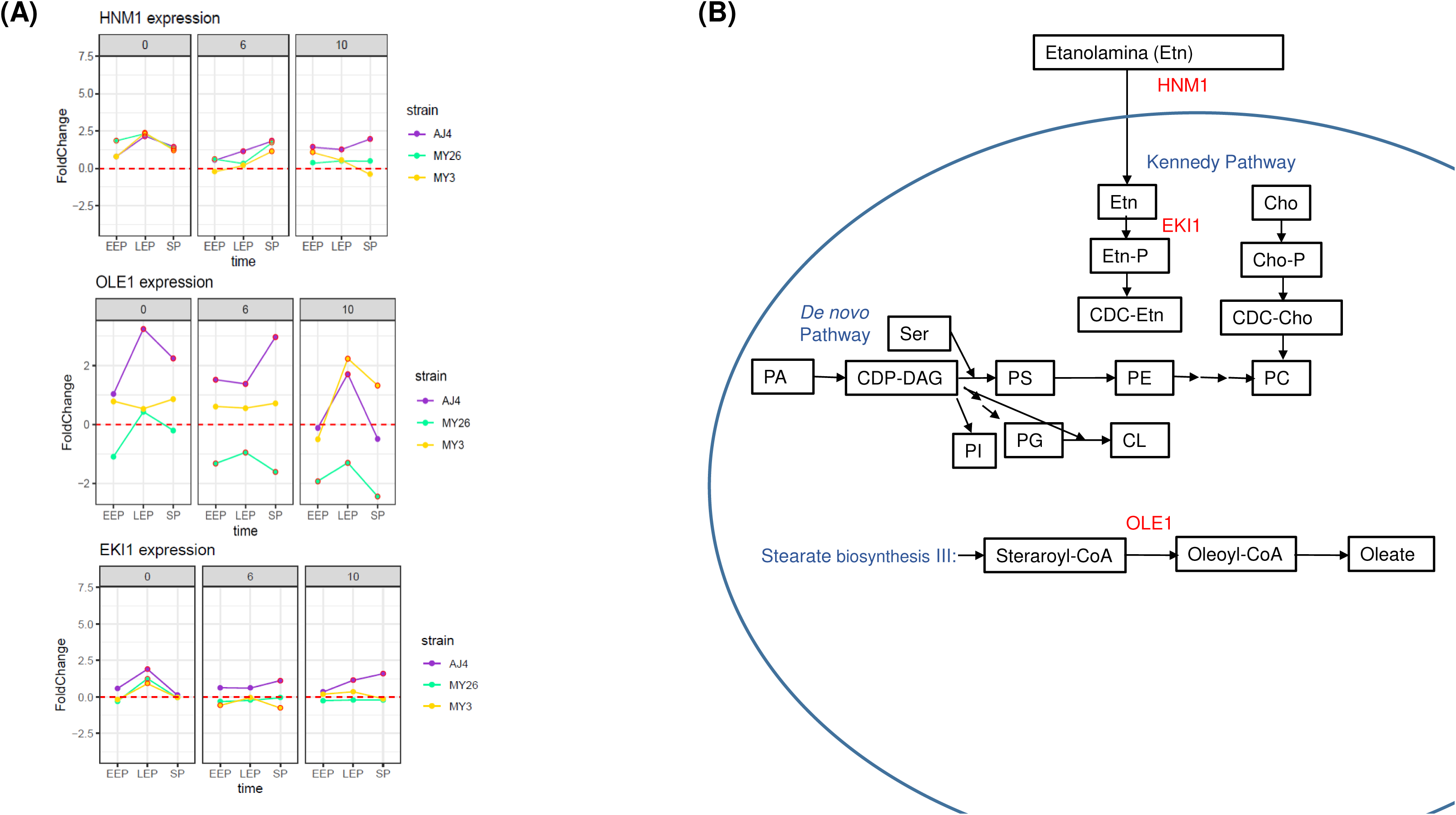
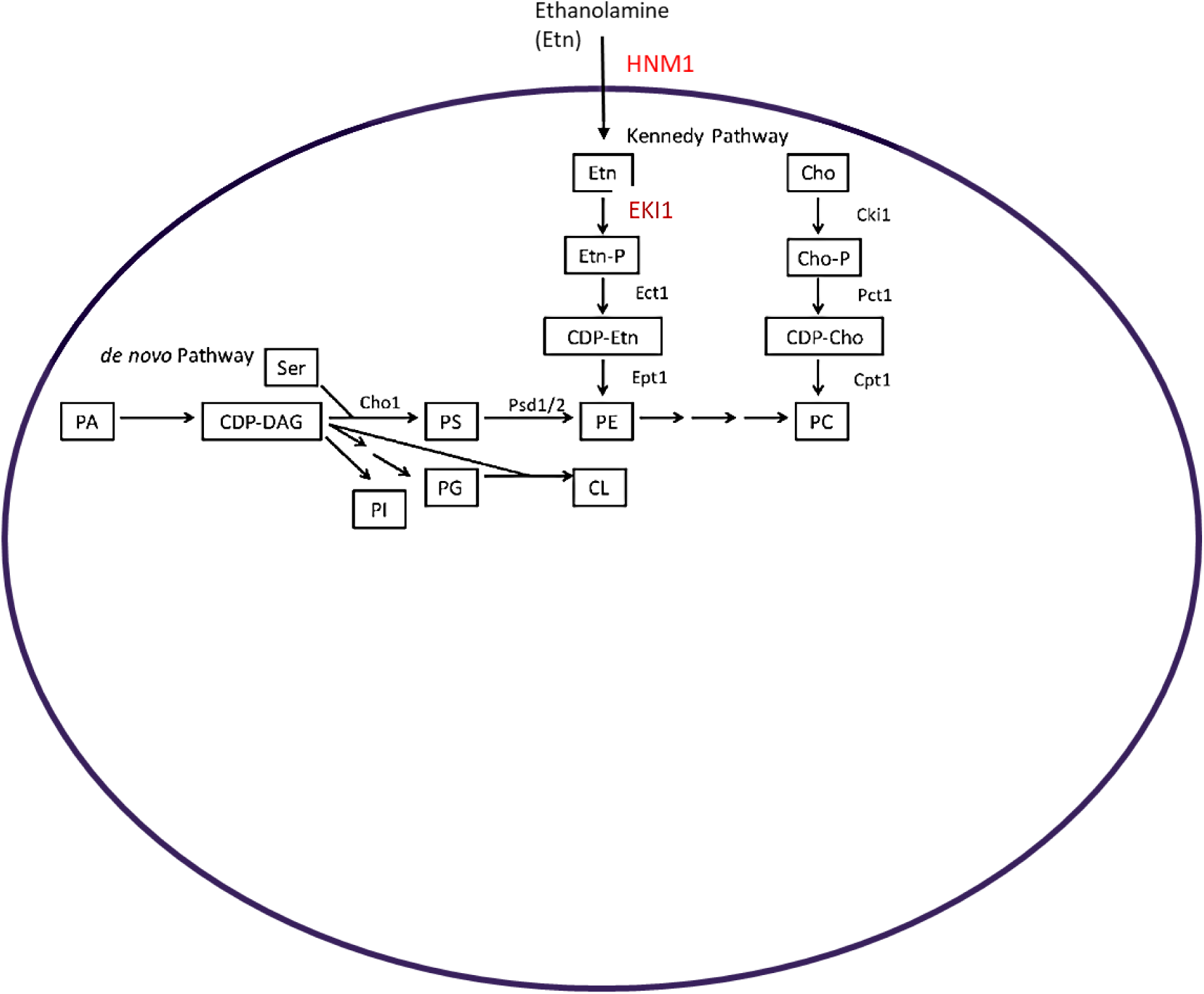
Results from the fold change analysis of ergosterol synthesis genes in *S. cerevisiae* strains AJ4, MY3, and MY26 were evaluated under three ethanol concentrations (0%, 6%, and 10%), and at three time points (*t1*, *t2*, *t3*), relative to the baseline expression level at *t0*, without ethanol supplementation.

Regarding the main membrane lipid biosynthesis processes, other genes exhibited differential behavior among strains. This includes *HMN1*, a transporter of phospholipid precursors such as choline and ethanolamine; and *EKI1*, the first enzyme in transforming ethanolamine into phosphatidylethanolamine. Both genes exhibited similar expression dynamics in the absence of ethanol for all three strains; however, their expression changed in the presence of ethanol, with AJ4 maintaining higher transcriptional levels for both genes (**Figure 4**). Additionally, *OLE1*, responsible for the desaturation step in the synthesis of oleic and palmitoleic acids, was repressed in MY26 under alcoholic conditions. In contrast, AJ4 and MY3 present higher expression levels (**Figure 4**).

**Figure 4.**
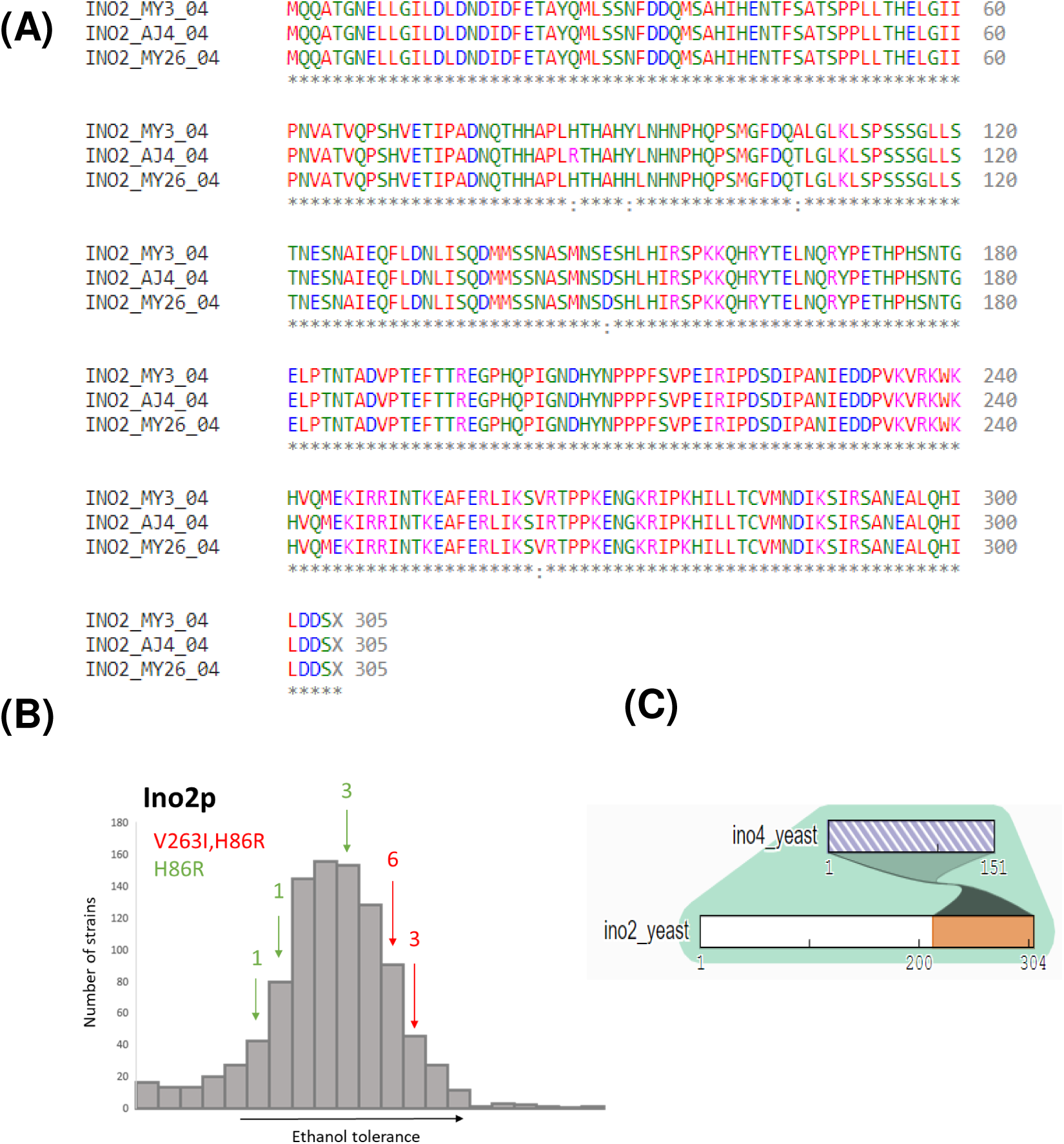
**(A)** Results of the expression analysis of genes *HMN1*, *EKI1*, and *OLE1* in *S. cerevisiae* strains AJ4, MY3, and MY26 were obtained under three ethanol concentrations (0%, 6%, and 10%), and at three-time points (*t1*, *t2*, *t3*), relative to the baseline expression level at *t0*, without ethanol supplementation. **(B)** A schematic representation of the phospholipid synthesis pathway involving the enzymes encoded by genes *OLE1*, *HMN1,* and *EKI1*.

### The role of the transcription factor Ino2p in the regulation of the response to ethanol stress

Remarkably, a transcriptional factor enrichment analysis made with *Yeastract*, using “DNA binding and expression evidence” (24), indicated that our set of differentially expressed genes is regulated by the transcription factor encoded by the gene *INO2* (Ino2p). A deeper investigation into *INO2* (*YDR123*) involved a comparative Ino2p protein sequence analysis for the selected strains. The result revealed two variants specific to AJ4 compared to the other two strains: a histidine-to-arginine substitution at position 86 (H86R) and a valine-to-isoleucine substitution at position 263 (V263I). Notably, Clustal Omega (25), a multiple sequence alignment tool, classified these amino acid replacements as conservative, indicating that they are likely to preserve the protein’s overall structure and function. Furthermore, we analyzed 979 Ino2p sequences from different strains, a highly representative set of *S. cerevisiae*’s diversity, including 1,011 strains from Peter et al. (26), to investigate the prevalence of the mutations observed in AJ4 across other strains. The V263I variant was identified in only nine strains, while the H86R variant was found in the same nine plus five additional strains. To explore the potential impact of these variants on ethanol tolerance, we referred to phenotypic data from Peter et al. (25), which included growth assays in media containing 15% ethanol. Strains exclusively harboring the H86R mutation exhibited relatively moderate ethanol tolerance. In contrast, strains carrying both mutations demonstrated noticeably higher ethanol tolerance levels, surpassing the median tolerance threshold (**Figure 5**). Thus, the AJ4 *INO2* allele could be a significant contributor to the strain’s higher ethanol tolerance.

**Figure 5.**
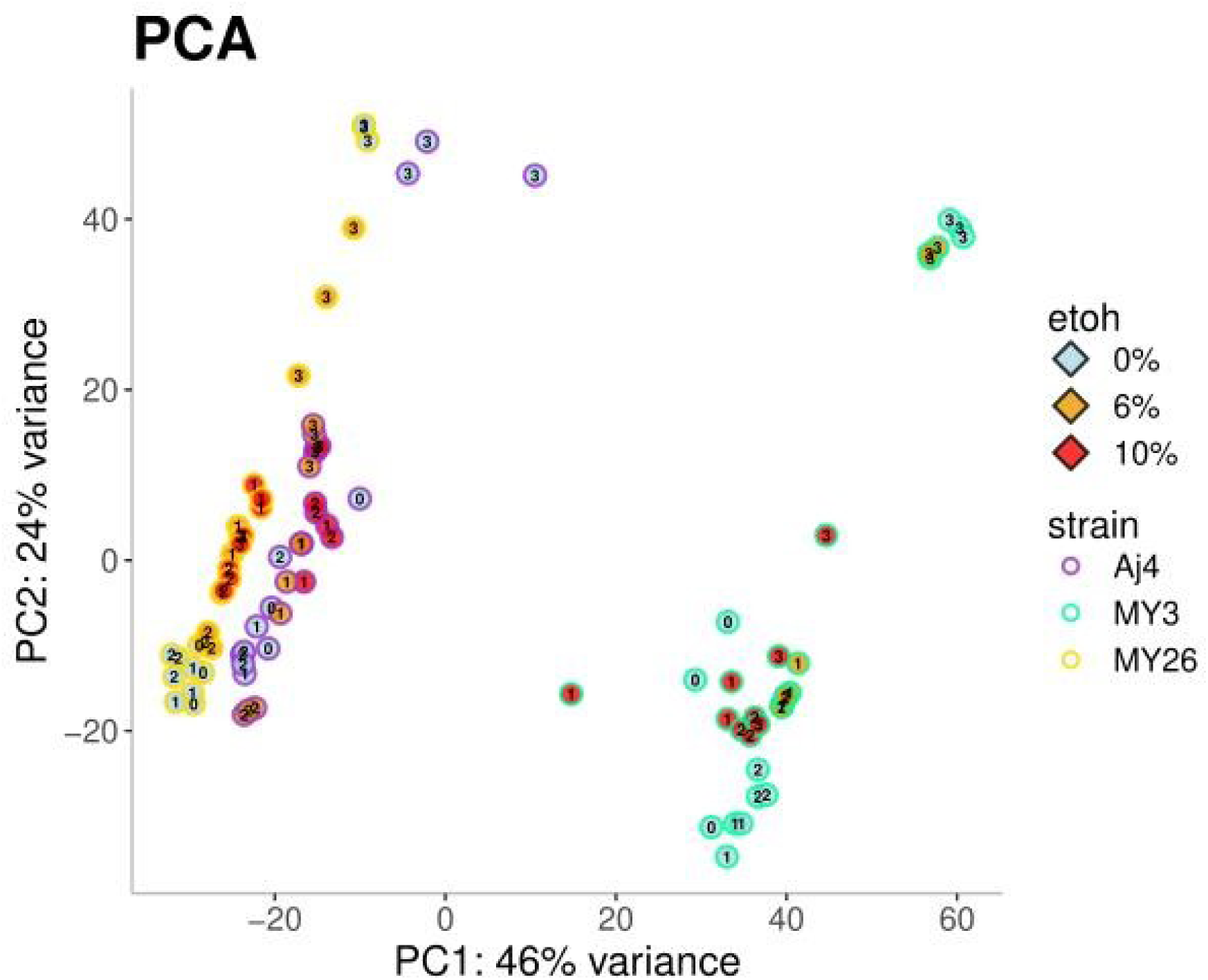
**(A)** Alignment of the amino acid sequence of Ino2p transcription factor from *S. cerevisiae* strains AJ4, MY3, and MY26. **(B)** Representation of the ethanol tolerance of 979 strains (Peter et al., 2018), indicating the presence of mutations in V263I and in H86R in Ino2p. **(C)** Representation of the interaction between Ino2p and Ino4p.

The transcriptional activator Ino2p exhibits a complex structure with distinct functional domains. The N-terminal region of Ino2p contains two crucial transcriptional activation domains: TDA1 and TDA2. TDA1 spans the first 35 amino acids of the protein sequence, while TDA2 is located between amino acids 101 and 135 (27). Additionally, Ino2p contains a Basic Helix-Loop-Helix (bHLH) domain, which interacts with Ino4p to bind specifically to the promoter DNA sequence by engaging with the major groove of the DNA. The bHLH domain in Ino2p is structured as follows: the basic region extends from amino acids 233 to 247, the first helix from 247 to 263, the loop from 263 to 276, and the second helix from 276 to 302, with the protein terminating at amino acid 304 (28). The interaction between Ino2p and Ino4p is crucial for the simultaneous binding to the promoter DNA fragment, as both proteins are required to form the functional complex (28).

The Ino2p^H86R^ variant features the amino acid replacement in a long loop within the N-terminal region of the protein. The substitution of histidine with arginine in a protein loop can have various effects depending on the specific location and the role of the residue in the protein’s structure and function. Differences in charge, size, flexibility, and hydrogen bonding potential between histidine and arginine may lead to changes in protein stability, folding, and interactions.

On the other hand, the AJ4 Ino2p^V263I^ variant has its amino acid replacement located at the beginning of the helix that interacts with Ino4p, near the binding site of the ligand, as observed in the predicted structure **(Figure 6)**. Although the replacement of valine to isoleucine is considered conservative, it can still affect protein structure, stability, and function depending on the specific location and context of the amino acid replacement. The additional methylene group in isoleucine may subtly alter the packing and hydrophobicity of the side chain, potentially leading to functional consequences. To investigate whether the Ino2p variants H86R and V263I impact *S. cerevisiae*’s ethanol tolerance, we conducted experimental tests.

**Figure 6.**
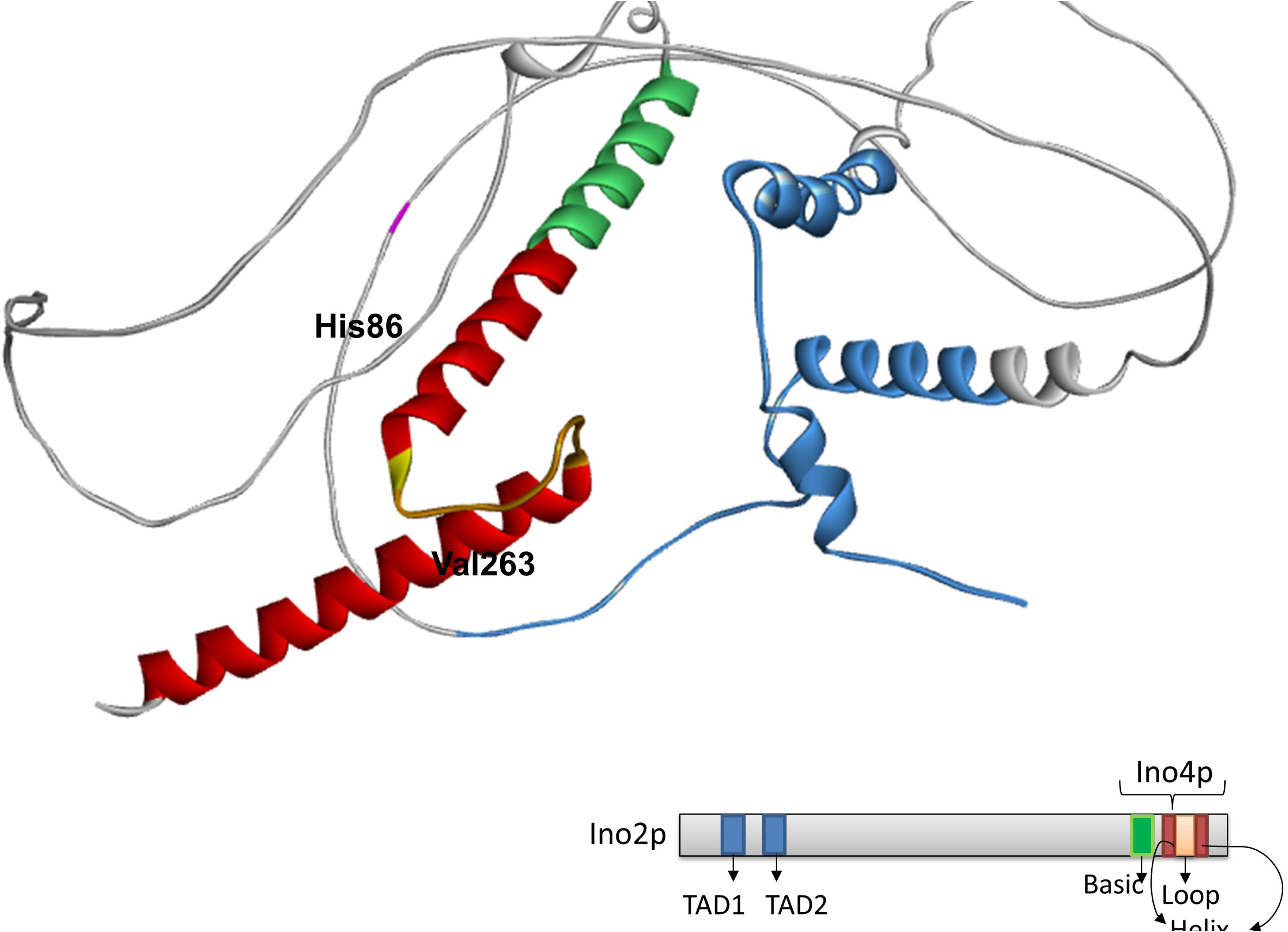
Structure of the transcription factor Ino2p. This structure was predicted using AlphaFold (53) and visualized with BIOVIA Discovery Studio (55). The TAD1 and TAD2 regions are depicted in blue; the basic domain where the bHLH begins is shown in green, and the helix of the domain is highlighted in red. Amino acids 86 and 263 are marked in pink and yellow, respectively.

### Ethanol tolerance in *INO2* mutants

To investigate whether V263I and H86R replacements could affect ethanol tolerance, we generated *INO2* mutants using CRISPR-Cas9 in the tolerant strain AJ4. The three mutants generated were: AJ4 (Ino2p^I263V^); AJ4 (Ino2p^R86H^); and AJ4 (Ino2^I263V, R86H^). Ethanol tolerance tests were then conducted at various concentrations to determine growth parameters. We calculated the areas under the growth curve (AUC) at different ethanol concentrations, and used these to determine the non-inhibitory concentrations (NIC) and maximum inhibitory concentrations (MIC) (**Figure 7**).

**Figure 7.**
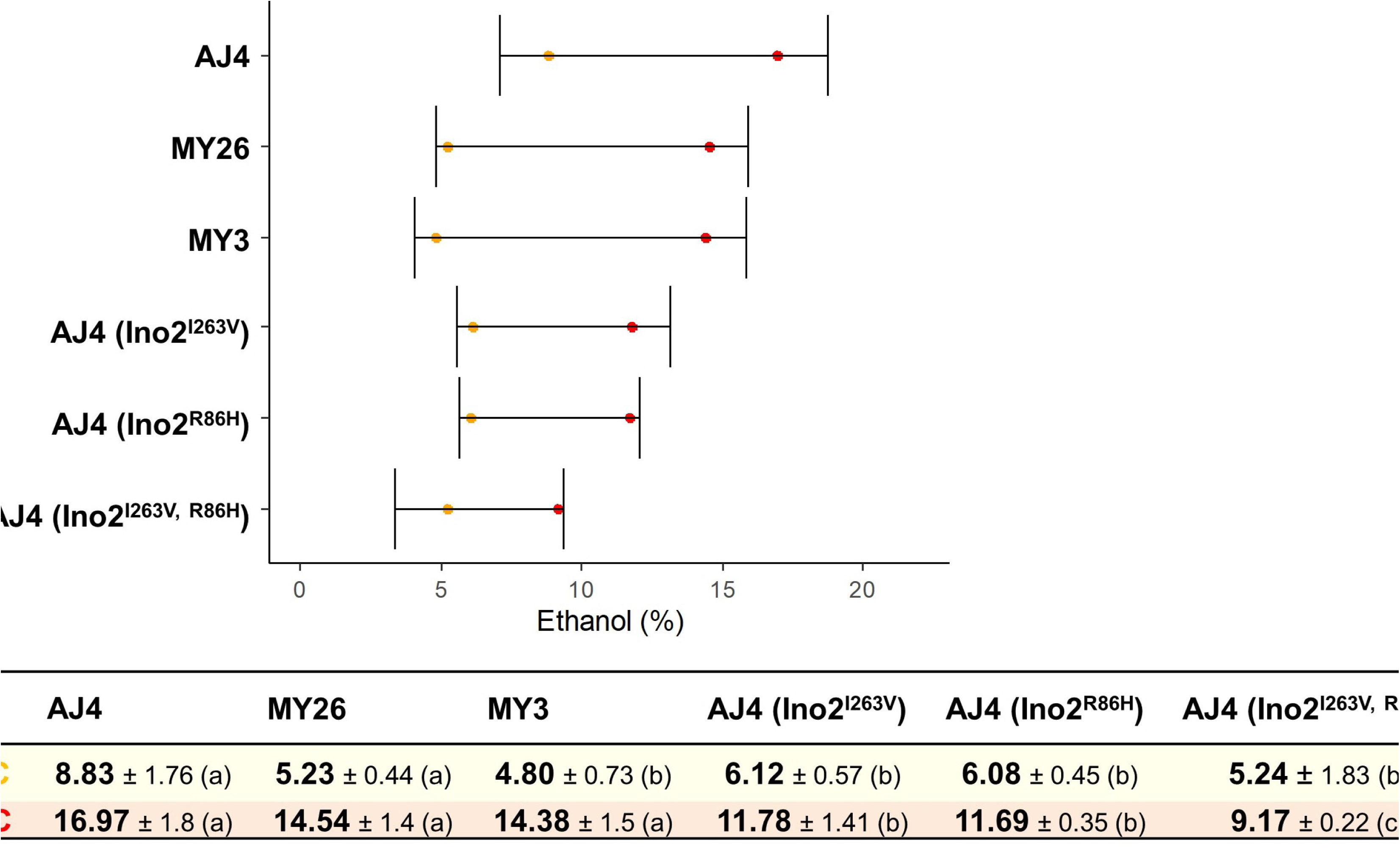
Ethanol non-inhibitory concentrations (NIC) and maximum inhibitory concentrations (MIC) of wild type AJ4, and the mutants AJ4 (Ino2p^I263V^), AJ4 (Ino2p^R86H^), and AJ4 (Ino2p^I263V, R86H^).

All mutants exhibited decreased NIC and MIC values, clearly indicating that both amino acid replacements contribute to ethanol tolerance in AJ4. The double mutant showed a more pronounced effect, with NIC values similar to those of the moderately and slightly tolerant strains, MY3 and MY26, and MIC values significantly lower. These findings suggest that, the presence of this *INO2* allele, encoding two amino acid replacements, could explain not only the differences in ethanol tolerance between AJ4 and the other two strains but also a more severe effect in the AJ4 genomic background.

In summary, the reversion of the AJ4 V263I and H86R replacements were observed to decrease ethanol tolerance, with a more pronounced effect at higher ethanol concentrations. This reduced tolerance may result from changes in the interaction between Ino2p and other transcription factors, such as Ino4p and Toa1p, which are involved in regulating genes related to membrane lipid biosynthesis.

## DISCUSSION

Crabtree-positive yeasts, particularly those of the *Saccharomyces* genus, have evolved high alcohol fermentation efficiency as part of their “make-accumulate-consume” ecological strategy (28). However, this also presents a significant challenge: the ethanol produced during fermentation is toxic to the yeast itself. To survive, *Saccharomyces* yeasts have developed various mechanisms to resist ethanol stress. Ethanol stress still represents a significant challenge for yeast cells in industrial biotechnology, underscoring the importance of elucidating the mechanisms involved in ethanol stress response. Elucidating these mechanisms is vital for developing targeted strategies to enhance yeast strains and optimize processes for improved sustainability (29). However, ethanol tolerance is a complex, strain-dependent trait (16), making it challenging to study.

In this study, we comprehensively analyzed gene expression under different ethanol concentration stress in three *S. cerevisiae* strains: the highly tolerant AJ4, the moderately tolerant MY3, and the slightly tolerant MY26. We observed some general stress responses, including mechanisms to avoid protein misfolding and aggregation, and to maintain cellular homeostasis (30). Additionally, we found that genes related to sporulation and mating were overexpressed, suggesting that yeast cells might enter a quiescence state, waiting for better conditions for vegetative growth, as reported in other studies on stress response (31). We also identified genes related to mitochondrial aerobic respiration, which is essential for energy generation (32) and maintaining cellular homeostasis under ethanol stress. Furthermore, sugar transporters, including *GAL2, HXT2, HXT3,* and *HXT4* (33) showed differential expression and have been previously linked to ethanol tolerance. (34).

Our main focus, however, was on the effect of ethanol stress on yeast membrane lipid metabolism genes, as changes in the membrane lipid composition is the first barrier against ethanol. Thus, genes involved in inositol monophosphate (IMP) synthesis were overexpressed in the tolerant strain AJ4, suggesting that maintaining lipid homeostasis is crucial in ethanol tolerance response (21). *ELO3*, which encodes a fatty acid elongase for synthesizing very long-chain fatty acids (20-26-carbons) from C18-CoA primers, was overexpressed in all three strains at the time point *t3*. Additionally, *ELO2,* encoding an elongase for fatty acids up to 26 carbons, was overexpressed in MY3 and MY26. Long-chain fatty acids are known to reduce membrane fluidity (35), a trait commonly associated with increased ethanol tolerance (21).

Ergosterol synthesis was the primary metabolic pathway repressed in MY3 and MY26 compared to AJ4, under ethanol stress. Ergosterol is the main sterol found in yeast cells, playing a critical role in regulating yeast membrane permeability and fluidity, particularly under high ethanol stress (36). By incorporating ergosterol into the lipid bilayer, it influences the packing and ordering of membrane lipids, thereby affecting the membrane’s physical properties. Traditionally, reducing membrane fluidity and permeability has been seen as a key strategy for ethanol tolerance (37–39). In our study, the strain-specific upregulation of ergosterol biosynthesis genes in the highly tolerant strain AJ4 aligns with its role in enhancing membrane compactness to achieve higher ethanol tolerance. However, previous research on *in vitro* membranes derived from AJ4 lipid extracts suggested that increased membrane fluidity contributes to a higher ethanol tolerance in AJ4 (21). These apparent contradiction could be due to the complexity of plasma membrane organization *in vivo*, where different subregions (rafts or islands) have variable compositions and, hence, different physical properties (40). One such ergosterol-rich subregion is associated to Can1p, a membrane arginine permease that requires phosphatidyl ethanolamine for its localization and is therefore exclusively found in lipid rafts. This specific association has been linked to increased tolerance to ethanol and other stresses (41). A higher ergosterol concentration in these subregions could be compatible with an overall increase in membrane fluidity elsewhere.

The transcription factor Ino2p plays a central role in regulating phospholipid biosynthesis genes in the yeast *S. cerevisiae* (42). The lipid biosynthesis genes share a common promoter region sequence (5’CATGTGAAAT3’), recognized as the inositol-sensitive upstream activation sequence (UAS)_INO_ (43, 44). The Ino2p-Ino4p complex binds to a fragment of the *INO1* promoter (45), containing the consensus-binding site for the bHLH family of proteins, also known as (UAS)_INO_ (28). This bHLH binding site is essential for the repression and activation of *INO1* and other phospholipid biosynthesis genes. *INO1* regulation is particularly important under ethanol stress, as upregulation of lipid metabolism-related genes like *INO1* is concurrent with the activation of the unfolded protein response (UPR) (13,46). Moreover, *INO1* is regulated by the TFIIA subunit Toa1p, which interacts physically with Ino2p and is necessary for the proper activation of genes containing the (UAS)_INO_ promoter element, especially *INO1*. Specifically, Ino2p binds to two separate structural domains of Toa1p, the N-terminal four-helix bundle structure, required for dimerization with Toa2p, and its C-terminal β-barrel domain, contacting TBP and sequences of the TATA element (27).

Interestingly, our sequence analysis of the Ino2p in the three strains, revealed that the highly ethanol-tolerant AJ4 strain harbors two amino acid replacements (V263I and H86R) compared to the less tolerant strains MY3 and MY26. We also identified these replacements in Ino2p sequences from strains with above-average ethanol tolerance (26). The V263I replacement impacts a helix in the bHLH involved in Ino2p-Ino4p interaction, potentially altering protein’s structure and function. Conversely, the H86R replacement is located in a loop domain of the N-terminal region and may affect binding with the TFIIA subunit Toa1p. Our study demonstrated that the reversion of these replacements in AJ4 led to a significant decrease in MIC for ethanol, suggesting that this Ino2p variant present in AJ4 plays a role in regulating genes involved in the ethanol tolerance response. Moreover, the AJ4-derived mutant with the double amino acid reversion exhibited lower ethanol tolerance than the less tolerant strains MY3 and MY26, indicating that the specific genetic background of each yeast strain can modulate with other mechanisms the effects of these genetic changes on the ethanol tolerance phenotype.

Ethanol tolerance is a complex trait involving multiple mechanisms, and lncRNAs act in a strain-specific manner in the ethanol stress response (17). This suggests that transcriptional regulators like Ino2p may modulate the expression and function of ethanol-responsive lncRNAs. The intricate interplay between trans-acting factors like Ino2p and the unique genomic background of each strain likely contributes to the complex balance of ethanol tolerance mechanisms.

Taken all together, *Saccharomyces* yeasts have evolved high-efficiency ethanol production as an ecological strategy to outcompete other microorganisms, as high ethanol levels are toxic to most organisms. However, ethanol also poses a stress to the yeast itself, damaging cellular structures and membranes. To survive, *Saccharomyces* yeasts have developed ethanol resistance mechanisms, including changes in membrane composition. In this study, we identified an allele of *INO2* that confers ethanol tolerance, encoding a variant of the transcription factor Ino2p, a key regulator of lipid biosynthesis. This discovery sheds light on the genetic regulation of lipid metabolism and membrane composition, offering potential advancements in biotechnological applications.

Our findings also highlight the challenge of predicting the effects of specific genomic variants, as their interactions with a strain’s unique genetic background can produce unpredictable outcomes. The interplay between trans-acting factors like Ino2p and a strain’s broader genomic context is crucial in determining ethanol tolerance. These results underscore the need to consider the entire genomic background when studying complex traits like ethanol tolerance. The evolutionary history and genetic variations in each strain create a finely tuned regulatory network that influences how specific mutations manifest, offering valuable insights for strain improvement and industrial applications.

## MATERIALS AND METHODS

### Media and strains

The diploid *Saccharomyces* strains AJ4, MY26, and MY3 have been used in this study due to their varying ethanol tolerances. In addition, their lipid compositions have been characterized by Lairón-Peris et al. (2021) (21). Strain AJ4 (Lallemand) is used for bioethanol production and exhibits high ethanol tolerance; MY3 (Lallemand), is a wine yeast used in the production of rosé and red wines, and has low tolerance to ethanol; MY2, a strain used in agave fermentation, is characterized by low to medium ethanol resistance. These strains were taken from 15% glycerol stocks and were maintained in GPY media.

### Yeast culture and sampling

Strains were inoculated in 500 mL GPY and grown overnight. From the preculture, 750 mL of GPY media were inoculated to an OD_600_ of 0.2 in 1L Erlenmeyer flasks. This was done in triplicates for each strain (AJ4, MY3, and MY26) and each condition (0%, 6%, and 10% of ethanol). The inoculated yeast cells were grown for 1 hour without stress to allow adjustment to the medium. Subsequently, ethanol was added up to reach the specified concentrations. Flasks were placed in an incubator at 28°C, with orbital agitation at 150 rpm. Aproximately 10^8^ cells were collected at different time points: before the ethanol addition (*t0*), early exponential phase (*t1*), late exponential phase (*t2*), and stationary phase (*t3*). Cells were harvested by centrifugation, frozen in liquid nitrogen, and stored at −80 °C.

### RNA-seq analysis

RNA isolation was performed following the phenol-chloroform method (47). Cells were washed with DEPC water, and then, treated with phenol tris, phenol-chloroform (5:1), and chloroform-isoamyl alcohol (24:1). The aquous phase was transferred to clean tubes, and nucleic acids were precipitated first with LiCl, and then with ethanol and sodium acetate. RNA was resuspended in RNase-free water. RNA concentration was measured using a Nanodrop, and RNA integrity was assessed with an Aligent Bioanalyzer. RNAseq libraries were prepared with the Illumina Truseq Stranded mRNA kit and sequenced on an Illumina Hiseq 2000, obtaining 75-bp paired-end reads. Reads were quality trimmed using Sickle (length 50, quality 23), aligned to the *S. cerevisiae* S288C reference genome using Bowtie2, and mapped reads were counted by Htseq-count (union mode) (48). Count data was imported, processed, and normalized by removing low-expressed genes and using the variance stabilizing transformation method in DESeq2 for sample clustering (49). Differential expression analysis was conducted with Limma package (v.3.32.2) (50). Counts were transformed to log cpm, and expression was adjusted by Limma-voom. GO term enrichment was performed with FunSpec (51), including differentially expressed genes with an adjusted P value < 0.05 (52). The information about the sequences area available in BioProject ID PRJNA1150416.

### Ino2p structure analysis

AlphaFold (53) was used to predict Ino2p structure, PDBe (54) for structural data, and BIOVIA Discovery Studio Visualizer (55) for structure and amino acids representation.

### Introducing point mutations in *INO2* by CRISPR-Cas9 system

The CRISPR-Cas System (56) was used in two steps for easier selection. First, we introduced kanMX in the regions surrounding codons for amino acids H86 or V263. KanMX was amplified from the plasmid pUG6A with designed primers with the *INO2* sequences (Additional information, Table S1) using PCR with Ex Taq Polymerase (Takara). We used the plasmid pRCC_N-Ino2, which contained the gRNA trageting *INO2* point mutations and other necessary components, including natMX6. This plasmid was created by amplifying pRCC_N using PCR with Phusion® polymerase (Thermo Fisher Scientific). Primers (Additional information, Table S1) contained the gRNA sequence designed with Geneious software. Then, strain AJ4 was transformed using the LiAc/SS Carrier DNA/PEG method (57), introducing the plasmid pRCC_N-Ino2, and the kanMX DNA fragment. Cells were plated in GPY with 300 ng/µL nourseothricin (clonNAT). After two days, colonies were replated on media with 300 ng/μL G418 to select kanMx-resistant colonies. These colonies were tested by PCR with custom-designed primers (Additional information, Table S1), to confirm kanMX replacement of *INO2*. Selected colonies were cleaned of the plasmid with ClonNAT resistance. In step two, we introduced the point mutation with CRISPR-Cas9 in the *INO2*-KanMX mutants. The fragment of DNA was designed by combining single sequences (forward and reverse) containing the desired mutation in the middle. The plasmid pRCC_N-kanMX-gRNA, containing the gRNA targeting kanMX, was used for transformation. Cells were plated on ClonNAT, and mutants were confirmed by PCRs with selected primers (Additional information, Table S1). The positive mutants were preserved in 15% glycerol at -80°C. Single mutants were used to create double mutants using the same procedure. Final mutants were AJ4 (Ino2p^I263V^), AJ4 (Ino2p^R86H^), and AJ4 (Ino2p^I263V, R86H^).

### Ethanol tolerance assay

Mutants AJ4 (Ino2p^I263V^), AJ4 (Ino2p^R86H^), and AJ4 (Ino2p^I263V, R86H^), and wild-type strains AJ4, MY26 and MY3 were cultured in 1 mL of GPY. After 16 hours, cultures were centrifuged, washed, and suspended in PBS. After 2 hours, a cell density was adjusted to 4 x 10^7^ cells/mL. Cells were inoculated into minimal medium YNB with different ethanol concentrations (0, 1, 3, 5, 7, 9, 11, 13, 15, and 17 %) in 96-well plates containing 220 µL of medium and 100 µL Vaseline oil (PharmpurR) to prevent ethanol evaporation.

A Stacker Microplate Handling System, attached to plate readers SPECTROstar Omega (BMG LABTECH), in a 25°C and 70% humidity chamber, measured OD_600_ to monitor growth. Microplates were shaken in orbital mode for 20 seconds each hour at 400 rpm. Yeast growth was analyzed using the Growth Curve Analysis Tool (GCAT) (58), and the area under the curve (AUC) was calculated. To assess the relative growth of yeast strains at different ethanol concentrations, we computed the fractional area (fa) by comparing the AUC at each ethanol concentration to that of the control condition.

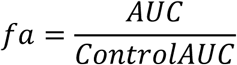

A modified Gompertz function was used to relate the fractional area (*y*) to the log of ethanol concentration (*x*) (59):

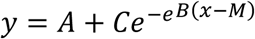

In this formula, A is the lower asymptote of *y*, B is the slope parameter, C is the distance between the upper and lower asymptote, and M is the log concentration of the inflection point. The values of the NIC and MIC are described as the intersection of the lines y = A+C and y = A, with the equation of the line tangential to the point (M) respectively.

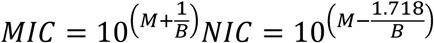

Eventually, the values of A, C, B, and M can be calculated using a non-linear fitting procedure, and NIC and MIC were determined.

## Authors’ contributions

**Sonia Albillos-Arenal**: Conceptualization, Investigation, Methodology, Validation, Formal analysis, Visualization, Writing – original draft.

**Javier Alonso del Real**: Conceptualization, Investigation, Methodology, Validation, Writing – original draft.

**María Lairón-Peris**: Investigation, Methodology.

**Eladio Barrio**: Conceptualization, Funding acquisition, Project administration, Supervision, Writing – review & editing

**Amparo Querol**: Conceptualization, Funding acquisition, Project administration, Supervision, Writing – review & editing

